# Genetic mapping of APP and amyloid-β biology modulation by trisomy 21

**DOI:** 10.1101/2022.03.10.483782

**Authors:** Paige Mumford, Justin Tosh, Silvia Anderle, Eleni Gkanatsiou Wikberg, Gloria Lau, Sue Noy, Karen Cleverley, Takashi Saito, Takaomi C Saido, Eugene Y. Yu, Gunnar Brinkmalm, Erik Portelius, Kaj Blennow, Henrik Zetterberg, Victor Tybulewicz, Elizabeth M.C. Fisher, Frances K. Wiseman

## Abstract

Individuals who have Down syndrome frequently develop early onset Alzheimer’s disease, a neurodegenerative condition caused by the build-up of aggregated amyloid-β and tau proteins in the brain. Amyloid-β is produced by *APP,* a gene located on chromosome 21. People who have Down syndrome have three copies of chromosome 21 and thus also an additional copy of *APP*; this genetic change drives the early development of Alzheimer’s disease in these individuals. Here we use a combination of next-generation mouse models of Down syndrome (Tc1, Dp3Tyb, Dp(10)2Yey and Dp(17)3Yey) and a knockin mouse model of amyloid-β accumulation (*App^NL-F^*) to determine how chromosome 21 genes other than *APP* modulate APP/amyloid-β in the brain when in three copies. We demonstrate that three copies of other chromosome 21 genes are sufficient to partially ameliorate amyloid-β accumulation in the brain. We go on to identify a subregion of chromosome 21 that contains the gene/genes causing this decrease in amyloid-β accumulation and investigate the role of two lead candidate genes *Dyrk1a* and *Bace2*. Thus an additional copy of chromosome 21 genes, other than *APP*, can modulate APP/amyloid-β in the brain under physiological conditions. This work provides critical mechanistic insight into the development of disease and an explanation for the typically later age of onset of dementia in people who have AD-DS compared to those who have familial AD caused by triplication of *APP*.

## Background

Down syndrome (DS) is caused by trisomy of human chromosome 21 (Hsa21), and occurs in around 1/1000 live births in Europe [1]. Most individuals who have DS develop the neuropathological features of Alzheimer’s disease; amyloid-β plaques and tau neurofibrillary tangles by the age of 50 [2], and 80% of individuals will have developed dementia by age 65 [3]. The high prevalence of AD in DS is in part due to the gene encoding amyloid precursor protein (*APP*) being located on Hsa21, thereby raising APP and amyloid-β protein levels [4–6]. Recent studies in preclinical systems have demonstrated that an extra copy of other genes on Hsa21 can modulate APP biology [7–10] and thus may alter the earliest stages of AD in individuals who have DS. These extra genes may act to *promote* or to *reduce* amyloid-β accummulation and which mechanism predominates is currently unclear. Notably the age of clinical dementia diganosis occurs slightly later in individuals who have DS compared with those who have early onset familial AD caused by duplication of *APP* (*i.e.*, with three copies of wild-type *APP*) [11]. However a direct comparison between these two causes of AD is confounded by the different diagnostic criteria used (as necessitated by the underlying intellectual disability that occurs in people who have DS) [12]. Understanding these processes is crucial to the appropriate selection of treatments for AD-primary prevention trials in people who have DS.

Previous *in vivo* studies have either examined the processing of endogenous mouse APP or used *APP* transgenic models to address this biology, but both of these approaches have limitations [7–9, 13]. Mouse APP differs in sequence from the human protein. In the amyloid-β region these differences both reduce the protein’s cleavage by β-secretase and the tendency of the amyloid-β generated to aggregate [14] thus limiting our ability to determine how changes to biology affect accumulation of amyloid-β – a key early aspect of AD. Whereas the over- and mis-expression of *APP* in transgenic mouse models may cause artefactual phenotypes, masking the modulatory effect of the extra copy of Hsa21 genes and causing elevated mortality which may confound data interpretation [13, 15].

Here we take a combinatorial approach: assessing the effect of an additional copy of Hsa21 genes (using a series of DS mouse models [16–18]) on the biology of endogenous mouse APP and on APP generated from a partially humanised *App* knock-in allele that does not cause elevated mortality (*App^NL-F^*) (Fig. 1) [15]. These data indicate that trisomy of genes on Hsa21 reduces amyloid-β accumulation and that people who have DS are partly protected from their raised *APP* gene dose by the additional copy of other genes on the chromosome. We go on to show that one of the gene or genes that cause this change in biology is located on mouse chromosome 16 between *Mir802* and *Zbtb21*. This region contains 38 genes and we specifically test if mechanisms linked to two lead candidate genes in this region, *Dyrk1a* and *Bace2*, occur in our novel *in vivo* AD-DS model system.

**Figure 1.**
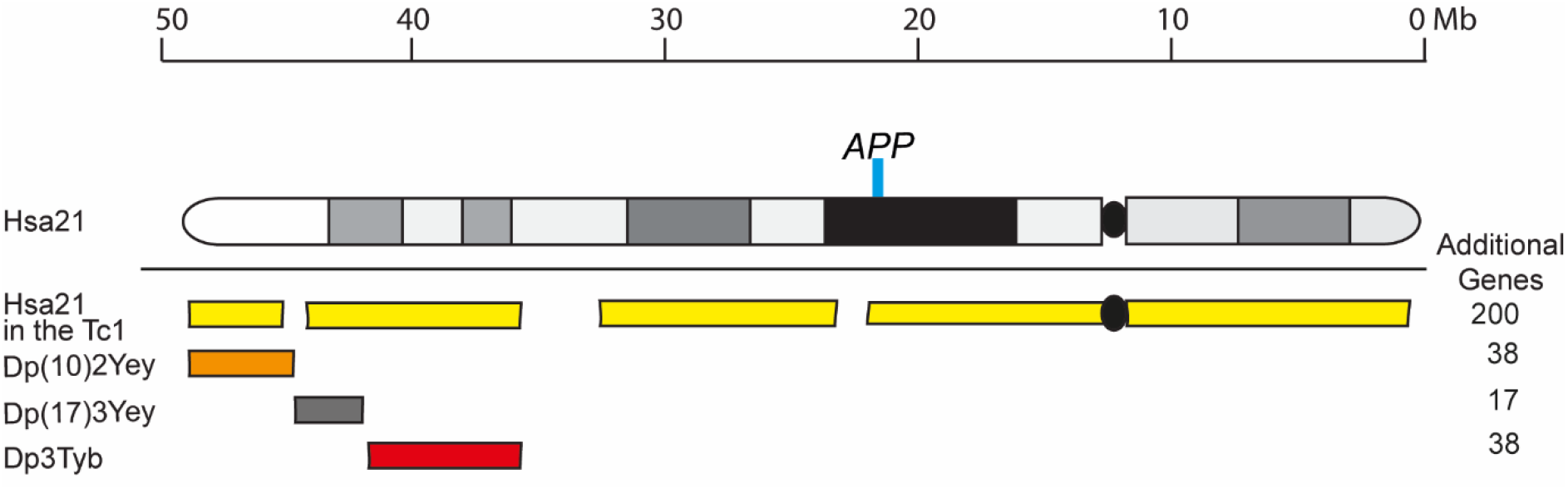
A schematic of Hsa21 indicating the major karyotypic bands, and the regions of Hsa21 or homologous regions of mouse chromosomes that are in three-copies in the mouse models used. The additional Hsa21 gene content in Tc1 (yellow), in the Dp3Tyb (red), Dp(10)2Yey (orange) and Dp(17)3Yey (grey) segmental duplication models of DS as indicated. The number of additional human genes in the Tc1, and the additional number of mouse genes in the Dp(10)2Yey, Dp(17)3Yey and Dp3Tyb as listed. The location of *APP/App* is indicated by the blue line. This colour scheme is used in subsequent figures to code for each of the mouse models.

## Methods

### Animal welfare and husbandry

All experiments were undertaken in accordance with the Animals (Scientific Procedures) Act 1986 (United Kingdom), after local institutional ethical review by the Medical Research Council, University College London and in accordance with ARRIVE2 guidelines [19].

Mice were housed in individually ventilated cages (Techniplast) with grade 5, autoclaved dust-free wood bedding, paper bedding and a translucent red “mouse house”. Free-access to food and water was provided. The animal facility was maintained at a constant temperature of 19-23°C with 55 ± 10% humidity in a 12 h light/dark cycle. Pups were weaned at 21 days and moved to standardised same-sex group housing with a maximum of 5 mice per cage.

The following mouse strains were used in this paper, here we show abbreviated name and then the official name and unique Mouse Genome Informatics (MGI) identifier: *App^NL-F^* (App^tm2.1Tcs^, MGI:5637816), Tc1 (Tc(HSA21)1TybEmcf, MGI:3814712), Dp3Tyb (Dp(16Mir802-Zbtb21)3TybEmcf, MGI:5703802), Dp(10)2Yey (Dp(10Prmt2-Pdxk)2Yey, MGI:4461400) and Dp(17)3Yey (Dp(17Abcg1-Rrp1b)3Yey, MGI:4461398).

Tc1 mice were maintained by mating Tc1 females to F1 (129S8 × C57BL/6) males. All other mouse strains were maintained by backcrossing males and females to C57BL/6J mice (imported from the Jackson Laboratory). Experimental cohorts for Tc1, Dp(10)2Yey and Dp(17)1Yey studies were produced by crossing mice carrying the additional Hsa21 or Hsa21 orthologous duplications with *App^NL-F/+^* animals in a two-generation cross to generate all required genotypes from the second generation (wild-type, Tc1/Dpx, *App^NL-F/NL-F^*, Tc1/Dpx;*App^NL-F/NL-F^*, Supplementary Fig. 1). As both the Dp3Tyb segmental duplication and the *App^NL-F^* gene are located on mouse chromosome 16 (Mmu16), for this cross we first generated a Dp3Tyb*-App^NL-F^* recombinant Mmu16, by crossing the two lines together and then back-crossing to C57BL/6J mice to identify recombined Mmu16, carrying both genetic changes on the same chromosome. Mice with the recombined Mmu16 were then crossed with *App^NL-F/+^* animals to generate Dp3Tyb*-App^NL-F/NL-F^* progeny. For this cross *App^NL-F/NL-F^* controls were generated from *App^NL-F/+^* x *App^NL-F/+^* matings, in addition to rare re-recombinations resulting in offspring without the Dp3Tyb segmental duplication but two copies of the *App* knock-in allele. Dp3Tyb controls were generated from Dp3Tyb x C57BL/6J mating’s generated at the same time as the Dp3Tyb*-App^NL-F/NL-F^* mice. Wild-type (WT) controls were taken from all three mating’s.

Animals were euthanized by exposure to rising carbon dioxide, followed by confirmation of death by dislocation of the neck in accordance with the Animals (Scientific Procedures) Act 1986 (United Kingdom).

### Tissue preparation and western blotting

For analysis of protein abundance in hippocampus and cortex, tissues were dissected under ice cold PBS before snap freezing. Samples were then homogenised in RIPA Buffer (150 mM sodium chloride, 50 mM Tris, 1% NP-40, 0.5% sodium deoxycholate, 0.1% sodium dodecyl sulfate) plus complete protease inhibitors (Calbiochem) by mechanical disruption. Total protein content was determined by Bradford assay or Pierce 660 nm assay (ThermoFisher). Samples from individual animals were analysed separately and were not pooled.

Equal amounts of total brain proteins were then denatured in LDS denaturing buffer (Invitrogen) and β-mercaptoethanol, prior to separation by SDS-PAGE gel electrophoresis using precast 4-12% Bis-Tris gels (Invitrogen). Proteins were transferred to nitrocellulose or PVDF membranes prior to blocking in 5% milk/PBST (0.05% Tween 20) or 5-10% bovine serum albumin (BSA)/PBST. Primary antibodies were diluted in 1% BSA/PBST, HRP-conjugated secondary anti-rabbit, anti-mouse and anti-goat antibodies (Dako) were diluted 1:10,000 in 1% BSA/PBST. Linearity of antibody binding was confirmed by a 2-fold dilution series of cortical protein samples. Band density was analysed using Image J. Relative signal of the antibody of interest compared to the internal loading control was then calculated, and relative signal was then normalized to mean relative signal of control samples electrophoresed on the same gel. Means of technical replicates were calculated and used for ANOVA, such that biological replicates were used as the experimental unit.

Primary antibodies against C-terminal APP (Sigma A8717, 1:10,000), β-actin (Sigma A5441, 1:60,000), DYRK1A (7D10, Abnova, 1:500) and BACE2 (Abcam ab5670 1:1,000) were used.

### Biochemical fractionation of mouse brain tissues for the analysis of human amyloid-β

Cortical proteins were fractionated as described in Shankar *et al.* (2009). A half cortex was weighed on a microscale and homogenised in 4 volumes of ice-cold Tris-buffered saline (TBS) (50mM Tris-HCl pH 8.0) containing a cocktail of protease and phosphatase inhibitors (Calbiochem) using a handheld mechanical homogeniser and disposable pestles (Anachem). Samples were then transferred to 1.5 ml microfuge tubes (Beckman Coulter #357448), balanced by adding more TBS and centrifuged at 175,000 × g with a RC-M120EX ultracentrifuge (Sorvall) fitted with rotor S100AT5 at 4 °C for 30 mins. Supernatant (the Tris-soluble fraction) was removed and stored at −80 °C. The remaining pellet was homogenised in 5 volumes of ice-cold 1% Triton-X100 (Sigma-Aldrich) in TBS (50mM Tris-HCl pH 8.0), balanced and centrifuged at 175,000 × g for 30 mins at 4 °C. The resultant supernatant (the Triton-soluble fraction) was removed and stored at −80 °C. The pellet was then re-suspended in 8 volumes (by original cortical weight) in TBS (50mM Tris-HCl pH 8.0), containing 5 M guanidine HCl and left overnight at 4 °C on a rocker to ensure full re-suspension, and subsequently stored at −80 °C. A Bradford assay or 660 nm protein assay (ThermoFisher) was performed to determine protein concentration.

### Biochemical preparation of mouse brain tissues for the analysis of mouse amyloid-β

A half cortex was weighed on a microscale and homogenised in 3 volumes of ice-cold Tris-buffered saline (TBS) (50mM Tris-HCl pH 8.0) containing a cocktail of protease and phosphatase inhibitors (Calbiochem) using a handheld mechanical homogeniser and disposable pestles (Anachem) based on the method in reference [20]. Homogenates were centrifuged at 21130 × g at 4 °C for 1 hour and the resultant supernatant was stored at −80 °C for onward analysis.

### Quantification of Aβ abundance by Meso Scale Discovery Assay

Amyloid-β_38_, amyloid-β_40_ and amyloid-β_42_ levels were quantified on Multi-Spot 96 well plates pre-coated with anti-amyloid-β_38_, amyloid-β_40_ and amyloid-β_42_ antibodies using multiplex MSD technology, as described in [9]. A 6E10 detection antibody was used to quantify human amyloid-β and 4G8 detection antibody for the quantification of mouse amyloid-β. Amounts of amyloid-β_38_, amyloid-β_40_ and amyloid-β_42_ were normalised to the original starting weight of cortical material.

### Immunohistochemistry

Half brains were immersion fixed in 10% buffered formal saline (Pioneer Research Chemicals) for a minimum of 48 hours prior to being processed to wax (Leica ASP300S tissue processor). The blocks were trimmed laterally from the midline by ~0.9-1.4 mm to give a sagittal section of the hippocampal formation. Two 4 μm sections at least 40 μm apart were analysed. The sections were pre-treated with 98% formic acid for 8 mins, followed by washing. The slides were wet loaded onto a Ventana XT for staining (Ventana Medical Systems, Tucson, AZ, USA). The protocol included the following steps: heat induced epitope retrieval (mCC1) for 30 mins in Tris Boric acid EDTA buffer (pH 9.0), superblock (8 mins) and manual application of 100 μl directly biotinylated mouse monoclonal IgG1 antibody against Aβ (82E1, IBL, 0.2 μg/ml) for 8 hours. Staining was visualised using a ChromoMap DAB kit followed by counterstaining with haematoxylin. The sections were dehydrated, cleared, and mounted in DPX prior to scanning (Leica SCN400F slide scanner). All images were analysed using ImageJ and by manual plaque counting by two independent researchers.

### Biochemical preparation of mouse brain tissues for mass spectrometry of amyloid-β

A half cortex was weighed on a microscale and was homogenized in 5 volumes of tris(hydroxymethyl)aminomethane (Tris)-buffered saline (TBS), pH 7.6, containing complete (cat: 04693116001, Roche) protease inhibitor. For the homogenization, one 5 mm bead per sample was used in a TissueLyser (Qiagen) for 4 min at 30 Hz. After homogenization, additional TBS with inhibitor was added up to 1mL and transferred to a new tube to be centrifuged at 31,000 *g* for 1 h at +4°C. The pellet was resuspended in 1 ml of 70% FA (v/v), followed by further homogenization in the TissueLyser for 2 min at 30 Hz and subsequent sonication for 30 s. The homogenate was centrifuged again at 31,000 *g* for 1 h at +4℃ and the supernatant (FA fraction) was dried down in a vacuum centrifuge.

Initially, 400 μl of 70% FA (v/v) added to the dried FA fractions, shaken for 30 min at 21°C and centrifuged at 31,000 *g* for 1 h at 4°C. After the removal of the supernatant, neutralization with 8 ml 0.5 M Tris was performed. IP performed as previously described with some modifications[21]. Briefly, 50 μl sheep anti-mouse magnetic beads (Thermo Fisher Scientific) that had previously been linked with 4 μg each of mouse monoclonal 6E10 and 4G8 (Biolegend) were added to the neutralized FA fraction. This complex was incubated overnight at +4℃ in 0.2% Triton X-100 in PBS (v/v). By using an automated magnetic-particle KingFisher ml system (Thermo Fisher Scientific), the samples were then washed with PBS Triton X-100, PBS, and 50 mM ammonium bicarbonate separately before elution in 100 μl 0.5% FA. Eluates were dried down in a vacuum centrifuge and stored at −80°C pending MS analysis.

### Mass spectrometry

Liquid chromatography-mass spectrometry (LC-MS) was conducted in a similar manner as described previously [22]]. Briefly, a nanoflow liquid chromatograph was coupled to an electrospray ionization (ESI) hybrid quadrupole–orbitrap tandem MS (Dionex Ultimate 3000 system and Q Exactive, both Thermo Fisher Scientific). Samples were reconstituted in 7 μl 8% FA/8% acetonitrile in water (*v/v/v*) and loaded onto an Acclaim PepMap 100 C18 trap column (length 20 mm; inner diameter 75 μm; particle size 3 μm; pore size 100 Å) for online desalting, and thereafter separated on a reversed-phase Acclaim PepMap RSLC column (length 150 mm, inner diameter 75 μm; particle size 2 μm; pore size 100 Å) (both Thermo Fisher Scientific). Mobile phases were A: 0.1% FA in water (*v/v*), and B: 0.1% FA/84% acetonitrile in water (*v/v/v*). The flow rate was 300 nl/min and a linear gradient of 3-40% B for 50 min was applied. The temperature of the column oven was 60 °C. Mass spectrometer setting were as follows: positive ion mode; mass-to-charge (*m/z*) interval 350-1800 *m/z* units; data dependent acquisition with 1 precursor ion acquisition (MS) followed by up to 5 fragment ion acquisitions (MS/MS); resolution setting 70,000 (for both MS and MS/MS); number of microscans 1 (MS and MS/MS); target values 10^6^ (MS and MS/MS); maximum injection time 250 ms (MS and MS/MS); fragmentation type was higher-energy collisional dissociation fragmentation (HCD); normalised collision energy (NCE) setting 25; singly charged ions and ions with unassigned charge were excluded for MS/MS selection.

Database search (including isotope and charge deconvolution) and label free quantification was performed with PEAKS Studio v8.5 (Bioinformatics Solutions Inc.) against a custom-made APP database. All suggested fragment mass spectra were evaluated manually.

MS signal was normalised to starting weight of cortex prior to data analysis.

### Statistical analysis and experimental design

All experiments and analyses were undertaken blind to genotype and sex. Experimental groups for all experiments were pseudo-randomised using 6-digit identification numbers allocated to each animal prior to genotyping. Data were analysed as indicated in figure legends by univariate ANOVA with between-subject factors being genetic status (*App^+/+^/App^NL-F/NL-F^* and/or wild-type/Hsa21 or wild-type/Dpx) and sex. Fractionation batch was included as a between-subject factor for analysis of amyloid-β by MSD assays. The subject means of technical replicates were calculated and used in the ANOVA for western blot and MSD assays, as the number of replicates for which data was available varied between samples. Repeat measures ANOVA was used for manual plaque counts (combining the data of two independent researchers).

## Results

### Trisomy of chromosome 21 decreases accumulation of amyloid-β in the cortex in the *App^NL-F^* mouse model

Following on from our previous studies, which indicated that an additional copy of Hsa21 alters APP biology and the accumulation of amyloid-β *in vivo* in an *APP* transgenic model [8, 9], here we determined if an additional copy of Hsa21 modulated APP/amyloid-β biology in the *App^NL-F^* knock-in mouse model. We undertook a two generation cross of the Tc1 mouse model of DS [18], which contains a freely segregating copy of Hsa21 (but not a functional additional copy of *APP*) [23], with the *App^NL-F^* model to generate 4 genotypes of mice (wild-type, Tc1, *App^NL-F/NL-F^* and Tc1;*App^NL-F/NL-F^*). We quantified the number of 82E1^+^ amyloid-β deposits in the cortex of these 4 genotypes of mice at 8-months of age. 82E1^+^ deposits were not observed in wild-type or Tc1 mice consistent with our previous study [9]. We found a significant decrease in the number of deposits in Tc1; *App^NL-F/NL-F^* compared with *App^NL-F/NL-F^* controls (Fig. 2 A, B).

**Figure 2:**
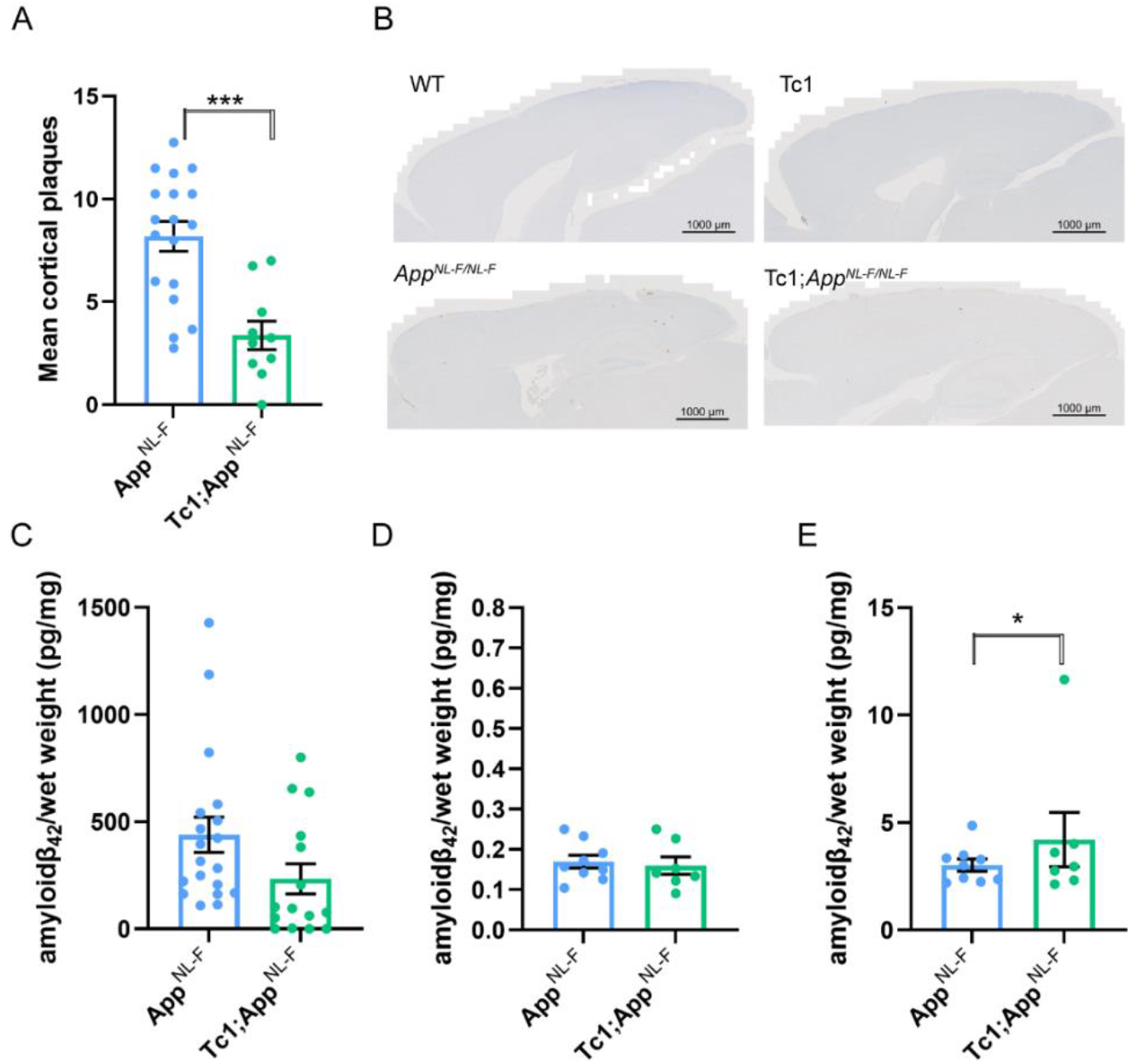
Trisomy of Hsa21 results in a decrease in amyloid-β deposition in the *App^NL-F/NL-F^* model. **(A-B)** Amyloid-β deposition (82E1) in the cortex was quantified at 8-months of age in male and female mice by manual plaque counting. **(A)** Significantly fewer amyloid-β deposits in the cortex were observed in Tc1;*App^NL-F/NL-F^* compared with *App^NL-F/NL-F^* controls (F(1,23) = 24.997, p < 0.001). *App^NL-F/NL-F^* female n=11, male n=7, Tc1; *App^NL-F/NL-F^* female n=4, male n=6; no amyloid-β deposits were observed in WT (female n=4) or Tc1 (female n=5, male n=3) age matched littermate controls (data not shown)**. (B)** Representative image of 82E1 stained amyloid-β deposits (brown) in wild-type (WT), Tc1, *App^NL-F/NL-F^*, Tc1;*App^NL-F/NL-F^* mice. **(C-E)** Total cortical proteins were biochemically fractionated and amyloid abundance analysed by MSD assay. **(C)** No statistically significant difference in the abundance of 5 M guanidine hydrochloride soluble amyloid-β_42_ was observed between Tc1;*App^NL-F/NL-F^* compared with *App^NL-F/NL-F^* controls (F(1,13) = 2.262, p = 0.156). *App^NL-F/NL-F^* female n=15, male n=8; Tc1*;App^NL-F/NL-F^* female n=8, male n=10. **(D)** No difference in the abundance of 1% Triton X-100 soluble amyloid-β42 was observed between Tc1;*App^NL-F/NL-F^* compared with *App^NL-F/NL-F^* controls (F(1,5) = 0.015, p = 0.907). *App^NL-F/NL-F^* female n=7, male n=2 (n = 8 below limit of detection); Tc1*;App^NL-F/NL-F^* female n=5, male n=2 (n = 11 below limit of detection). **(E)** Significantly more Tris soluble amyloid-β_42_ was observed in Tc1; *App^NL-F/NL-F^* compared with *App^NL-F/NL-F^* controls (F(1,5) = 10.697, p = 0.022). *App^NL-F/NL-F^* female n=7, male n=2 (n = 8 below limit of detection) Tc1*;App^NL-F/NL-F^* female n=5, male n=2 (n = 11 below limit of detection). Error bars show SEM, data points are independent mice.

We also determined if trisomy of Hsa21 modulated the biochemical aggregation of amyloid-β_40_ and amyloid-β_42_ in the cortex at 8-months of age, using biochemical protein fractionation by step-wise homogenisation and ultracentrifugation in sequentially more disruptive solutions (Tris-HCl, Tris-HCl 1% triton and finally 5 M guanidine hydrochloride). We then quantified amyloid-β_40_ and amyloid-β_42_ in each fraction normalised to starting brain weight (6E10 MSD triplex assay). Amyloid-β_42_ in the guanidine hydrochloride fraction was not significantly reduced in the presence of the extra copy of Hsa21 (Fig. 2C). A significant increase in Tris soluble amyloid-β_42_ was seen in the cortex of Tc1;*App^NL-F/NL-F^* compared with *App^NL-F/NL-F^* controls (Fig. 2E). No significant difference in Triton soluble amyloid-β_42_ abundance was observed (Fig. 2D). The amount of human amyloid-β_40_ in the *App^NL-F/NL-F^* model is very low because of the Iberian mutation in the modified *App* allele and this analyte was below the limit of detection in the Tris soluble fraction and did not significantly differ between genotypes in the Triton and 5 M guanidine hydrochloride fractions (Supplementary Fig. 2). These data indicate that an additional copy of a Hsa21 gene or genes is sufficient to reduce the deposition of amyloid-β in the cortex.

### Decreased amyloid-β accumulation in the Tc1-***App^NL-F/NL-F^*** model does not occur because of a reduction of APP or CTF-β abundance.

The significant increase in Tris soluble amyloid-β_42_ in the Tc1;*App^NL-F/NL-F^* cortex suggests that the decrease in amyloid-β accumulation observed is likely to be caused by a change to peptide aggregation. However, previous studies have suggested that genes on Hsa21 other than *APP* can increase APP protein level *in vivo* and modulate the abundance of the amyloid-β precursor APP-C-terminal fragment-β (CTF-β) [7, 9, 24]. Thus, we used western blotting to determine if the additional Hsa21 altered APP or CTF-β abundance in our new model system. Here, we found no evidence of decreased mouse or human full-length APP (FL-APP) in the cortex of the Tc1 and Tc1;*App^NL-F/NL-F^* compared with wild-type and *App^NL-F/NL-F^* mice (Fig. 3A, D). We note that significantly less FL-APP is detected in humanised models compared with controls (using antibody A8717), this may reflect a reduction in antibody binding rather than a biological reduction in protein level.

**Figure 3:**
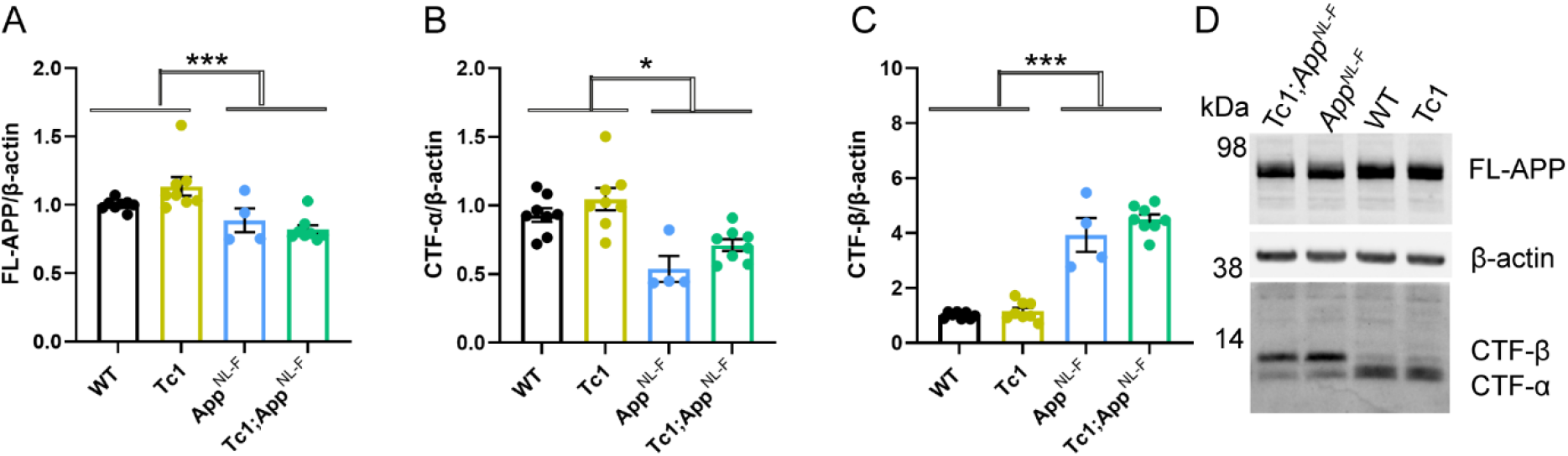
The abundance of FL-APP and CTF is not altered by trisomy of Hsa21. **(A-D)** The relative abundance of full-length APP (FL-APP), APP β-C-terminal fragment (β-CTF) and APP α-C-terminal fragment (α-CTF) compared to β-actin was measured by western blot using A8717 primary antibody in the cortex at 3-months of age in female and male mice. **(A)** Significantly less FL-APP was observed in mice in which *App* was humanised and mutated (F(1,19) = 23.837, p < 0.001). An additional copy of Hsa21 did not alter FL-APP abundance (F(1,19) = 0.599, p = 0.449). **(B)** Significantly less CTF-α was observed in mice in which *App* was humanised and mutated (F(1,19) = 5.950, p = 0.025) but an additional copy of Hsa21 did not alter α-CTF abundance (F(1,19) = 3.012, p = 0.099). **(C)** Significantly more β-CTF was observed in mice in which *App* was humanised and mutated (F(1,19) = 868.431, p < 0.001). By ANOVA a significant effect of Hsa21 on CTF-β abundance was detected (F(1,19) = 23.462, p < 0.001), however wild-type (WT) and Tc1 (Bonferroni pair-wise comparison p = 1.000) and *App^NL-F/NL-F^* and Tc1;*App^NL-F/NL-F^* (Bonferroni pair-wise comparison p = 0.118) were not statistically significant. WT female n= 4, male = 4: Tc1 female n = 3, male = 5; *App^NL-F/NL-F^* female n = 2, male n = 2; Tc1*;App^NL-F/NL-F^* female n = 4, male n = 4. **(D)** Representative image of western blot in WT, Tc1, *App^NL-F/NL-F^*, Tc1;*App^NL-F/NL-F^* mice. Error bars show SEM, data points are independent mice.

We found a signficant increase in CTF-β and a significant decrease in CTF-α in the cortex of *App^NL-F/NL-F^* mice consistent with the reported effects of the introduced mutations on APP processing (Fig. 3B, C, D) [15]. An additional copy of Hsa21 did not significantly alter wild-type APP CTF-β or CTF-α levels in the cortex of the Tc1 compared to wild-type-mice or in the Tc1;*App^NL-F/NL-F^* compared to *App^NL-F/NL-F^*. This finding contrasted with the large increase in CTF-β in male Tc1;*APP* transgenic model that we previously reported [9]. These data suggest that the reduction in deposition of amyloid-β in the Tc1;*App^NL-F/NL-F^* model is likely mediated by an enhancement of amyloid-β clearance or an impairment of peptide aggregation rather than a decrease in APP or CTF-β abundance.

### Decreased accumulation of amyloid-β is caused by an additional copy of 38 Hsa21 orthologous genes

Many DS-associated phenotypes are multigenic, caused by the combined effect of multiple Hsa21 genes acting together on one biological pathway. To understand the mechanisms underlying the decrease in amyloid-β accumulation in the Tc1;*App^NL-F/NL-F^* model further, we used a series of mouse models of DS that carry an extra-copy of subregions of mouse chromosomes that are orthogolous with Hsa21, to identify the combination of regions/genes responsible for the changes (Fig. 1) [16, 17]. We used 3 mouse lines that carry similar gene content to the Tc1 mouse model. However, because of limitations of available models we were not able to explore the Hsa21 genes closest to *App* on Mmu16 that are in three copies in the Tc1 model; as using recombination to generate the required combination of alleles in the available Dp9Tyb model was not feasible.

An extra copy of the genes between *Mir802* and *Zbtb21* (Dp3Tyb) was sufficient to decrease the accumulation of amyloid-β in the cortex, as quantified by 82E1 plaque counts (Fig. 4A). However, an extra copy of the genes between *Prmt2* and *Pdxk* (Dp(17)3Yey) and between *Abcg1* and *Rrp1b* (Dp(10)2Yey) was not sufficient to significantly alter the accumulation of amyloid-β in the cortex, as quantified by 82E1 plaque counts in 8-month old mice (Fig. 4B, C).

**Figure 4:**
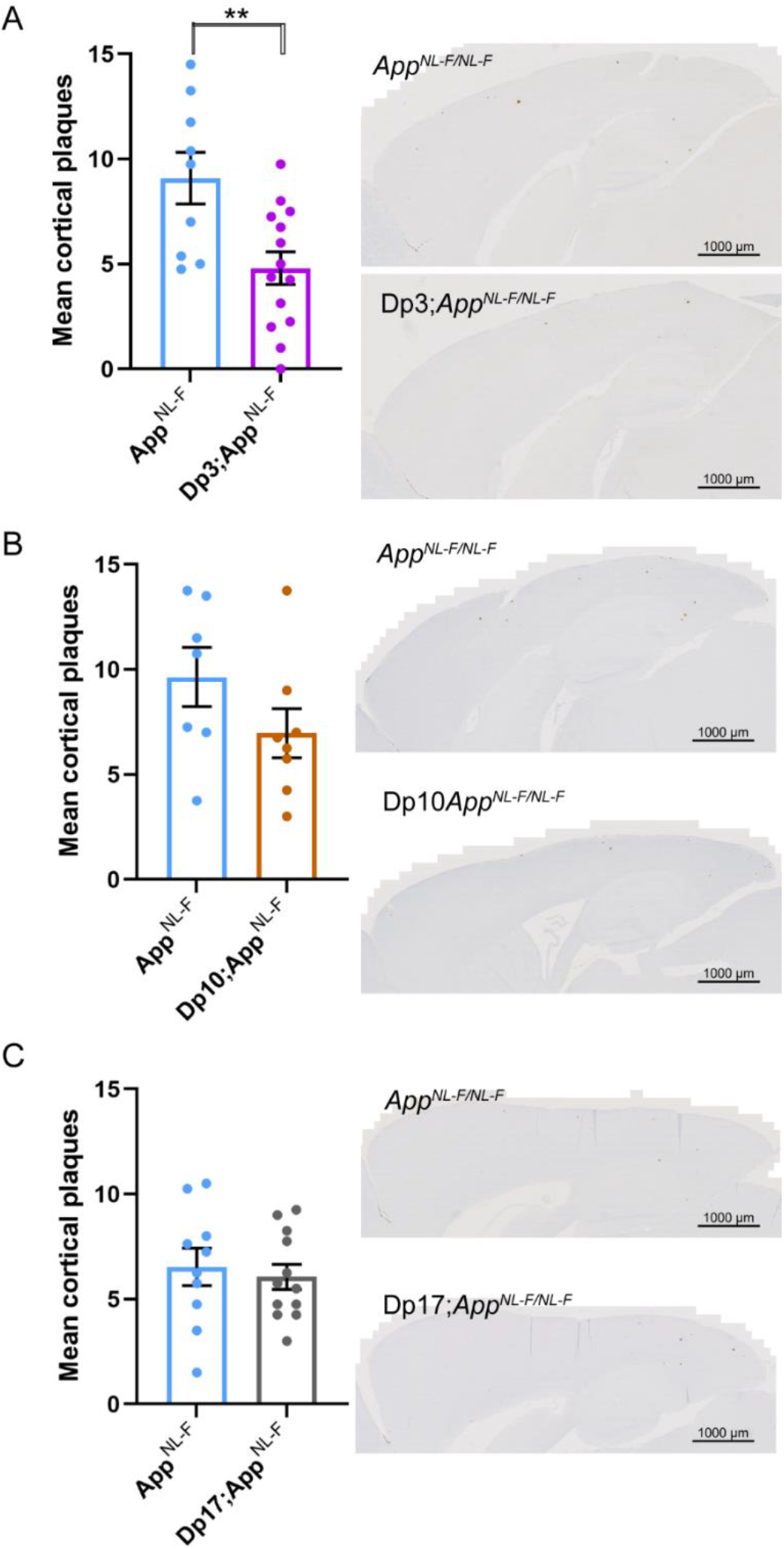
Duplication of the Dp3Tyb region of mouse chromosome 21 orthologous with Hsa21 is sufficent to decrease amyloid-β in the *App^NL-F/NL-F^* model. **(A-C)** Amyloid-β deposition (82E1) in the cortex was quantified at 8-months of age in male and female mice by manual plaque counting. **(A)** Significantly fewer amyloid-β deposits in the cortex were observed in Dp3Tyb;*App^NL-F/NL-F^* compared with *App^NL-F/NL-F^* controls (F(1,18) = 12.359, p = 0.002). *App^NL-F/NL-F^* female n= 2, male n= 7 Dp3Tyb;*App^NL-F/NL-F^* female n=8, male n=5. **(B)** A non-significant trend for reduced amyloid-β deposits in the cortex was observed in Dp(10)2Yey; *App^NL-F/NL-F^* compared with *App^NL-F/NL-F^* controls (F(1,11) = 12.359, p = 0.077). *App^NL-F/NL-F^* female n = 4, male n = 3 Dp10;*App^NL-F/NL-F^* female n = 6, male n = 2. **(C)** No difference in amyloid-β deposits in the cortex were observed in Dp(17)3Yey;*App^NL-F/NL-F^* compared with *App^NL-F/NL-F^* controls (F(1,18) = 0.021, p = 0.885). *App^NL-F/NL-F^* female n= 6, male n= 4 Dp(17)Yey *App^NL-F/NL-F^* female n= 6, male n= 6. Dp3Tyb abbreviated to Dp3, Dp(10)2Yey abbreviated to Dp10, Dp(17)3Yey abbreviated to Dp17 for clarity. Error bars show SEM, data points are independent mice.

The abundance of amyloid-β42 in the guanidine hydrochloride fraction was not significantly altered in the Dp3Tyb;*App^NL-F/NL-F^*, Dp(10)2Yey;*App^NL-F/NL-F^* or Dp(17)3Yey;*App^NL-F/NL-F^* mice compared with *App^NL-F/NL-F^* controls at 8-months of age (Fig. 5A-C). No significant difference in the abundance of Tris or Triton soluble amyloid-β42 was observed in Dp3Tyb;*App^NL-F/NL-F^*, Dp(10)2Yey;*App^NL-F/NL-F^* or Dp(17)3Yey;*App^NL-F/NL-F^* compared to controls (Fig. 5D-I). No difference in the abundance of guanidine hydrochloride or Triton soluble amyloid-β40 was observed between Dp(10)2Yey;*App^NL-F/NL-F^* or Dp(17)3Yey;*App^NL-F/NL-F^* compared to controls (Supplementary Fig. 3). These data indicate that a gene or genes in 3-copies in the Dp3Tyb model is sufficient to decrease the deposition of amyloid-β in the brain.

**Figure 5:**
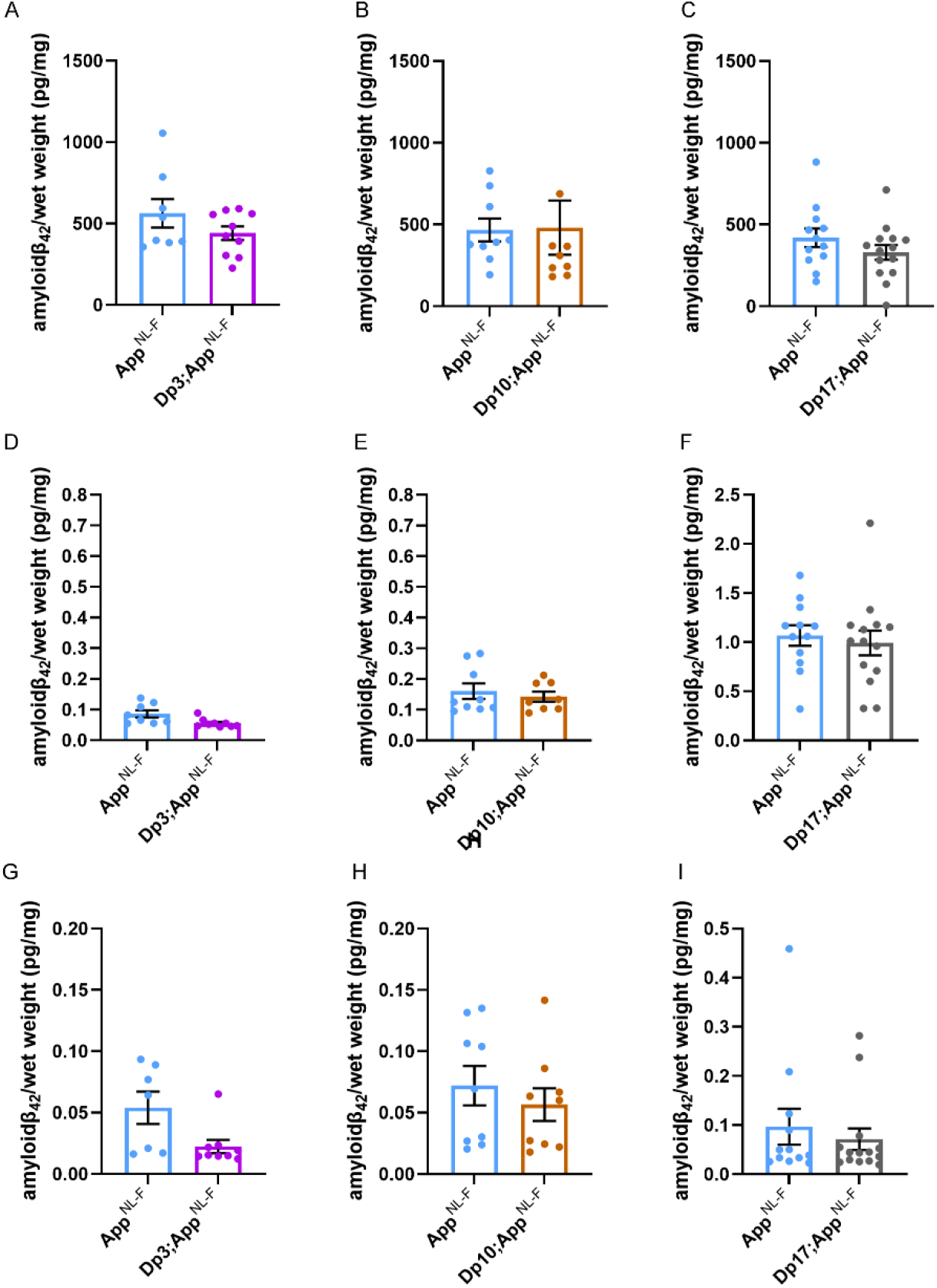
Biochemical solubility of amyloid-β is not altered by an additional copy of the Dp3Tyb, Dp(10)2Yey or Dp(17)Yey Hsa21 orthologous regions. Total cortical proteins were biochemically fractionated from 8-month of age mice and amyloid-β abundance analysed by MSD assay (6E10). No difference in the abundance of **(A)** 5 M guanidine hydrochloride soluble amyloid-β_42_ (F(1,12) = 0.128, p = 0.726), **(D)** 1% Triton X-100 soluble amyloid-β_42_ (F(1,12) = 2.863, p = 0.116) or **(G)** Tris soluble amyloid-β_42_ (F(1,10) = 0.281, p = 0.608) was observed between Dp3Tyb;*App^NL-F/NL-F^* compared with *App^NL-F/NL-F^* controls. *App^NL-F/NL-F^* female n = 2, male n = 6; Dp3Tyb*;App^NL-F/NL-F^* female n = 7, male n = 3. No difference in the abundance of **(B)** 5 M guanidine hydrochloride soluble (F(1,11) = 2.337, p = 0.115), **(E)** 1% Triton X-100 soluble amyloid-β_42_ (F(1,11) = 0.145, p = 0.711) or **(H)** Tris soluble amyloid-β_42_ (F(1,11) = 0.001, p = 0.978) was observed between Dp(10)2Yey;*App^NL-F/NL-F^* compared with *App^NL-F/NL-F^* controls. *App^NL-F/NL-F^* female n = 4, male n = 5; Dp3Tyb*;App^NL-F/NL-F^* female n = 6, male n = 3. No difference in the abundance of **(C)** 5 M guanidine hydrochloride soluble (F(1,20) = 4.079, p = 0.057), **(F)** 1% Triton X-100 soluble amyloid-β_42_ (F(1,20) = 0.124, p = 0.728) or **(I)** Tris soluble (F(1,20) = 0.521, p = 0.479) amyloid-β_42_ was observed between Dp(17)3Yey;*App^NL-F/NL-F^* compared with *App^NL-F/NL-F^* controls. *App^NL-F/NL-F^* female n = 7, male n = 5; Dp(17)Yey*;App^NL-F/NL-F^* female n = 7, male n = 7. Dp3Tyb abbreviated to Dp3, Dp(10)2Yey abbreviated to Dp10, Dp(17)3Yey abbreviated to Dp17 for clarity. Error bars show SEM, data points are independent mice.

### Increased DYRK1A does not lead to increased APP or amyloid-β in the Dp3Tyb model of DS

The Dp3Tyb model contains an additional copy of 38 genes, including *Dyrk1a*, a kinase which phosphorylates APP [25], increasing the abundance of the protein *in vivo*, contributing to raised soluble amyloid-β abundance in the Ts65Dn mouse model of DS [7]. We wanted to determine whether we could observe this previously reported biology in our new model system. We found that an extra copy of the Dp3Tyb region raises the abundance of DYRK1A in the cortex of 3-month old animals, including in the context of *App^NL-F^* knock-in mutations (Fig. 6A-B). This is consistent with numerous previous reports of dosage sensitivity of DYRK1A in the mouse throughout lifespan [7, 26, 27]. We also note that, consistent with previous reports in other mouse model systems, 3-copies of *Dyrk1a* in the Dp3Tyb*;App^NL-F/NL-F^* model was associated with an increase in cortical weight [28] (Supplementary Fig. 4A).

**Figure 6:**
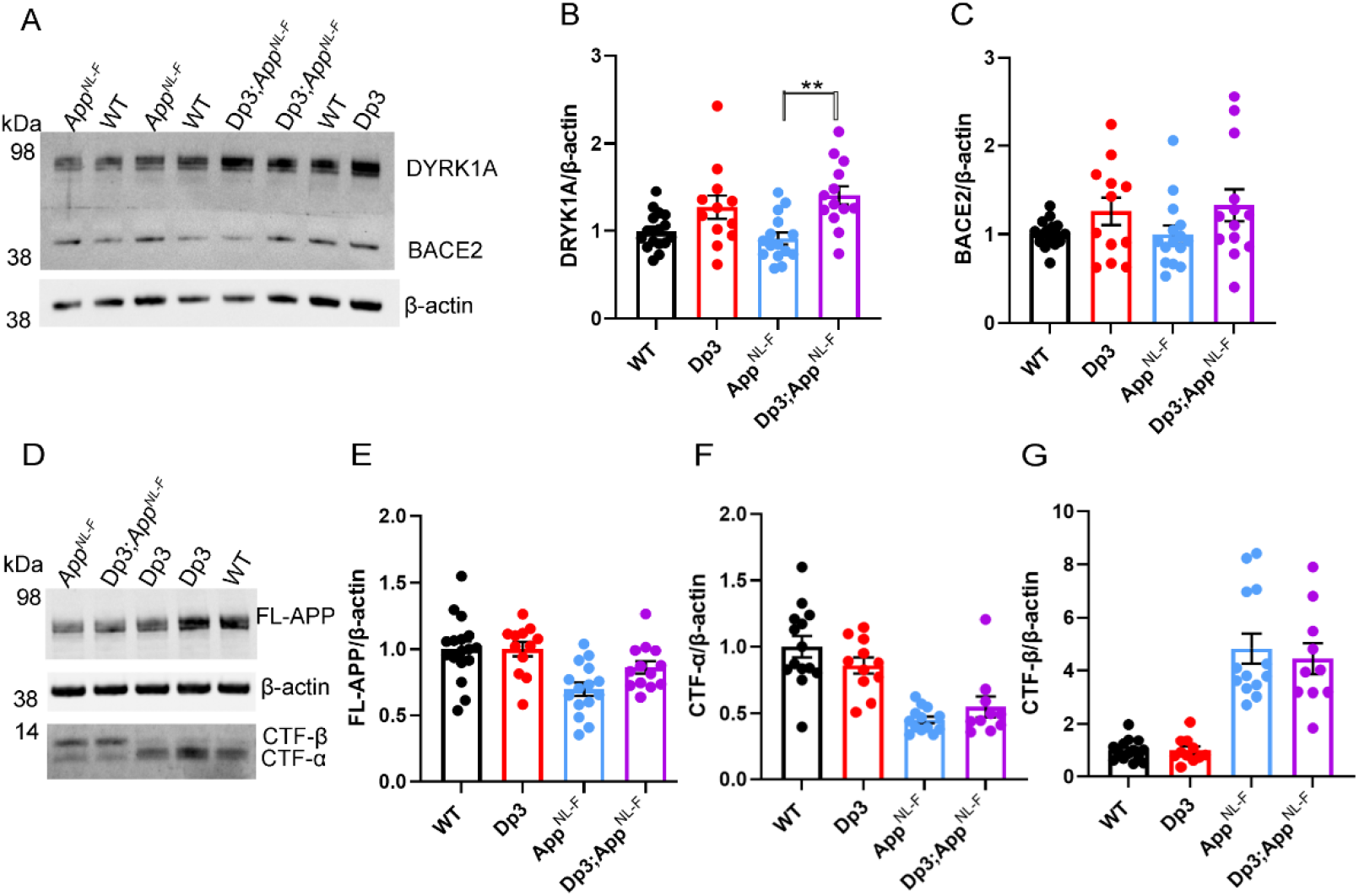
Duplication of Dp3Tyb region is sufficent to raise the protein abundance of DYRK1A in the cortex but does not significantly alter BACE2, FL-APP or CTF abundance. The abundance of **(A-B)** DYRK1A, **(A, C)** BACE2, **(D, E)** full-length, APP (FL-APP), **(D, F)** C-terminal fragment-α (CTF-α) and **(D, G)** C-terminal fragment-β (CTF-β) relative to β-actin loading control was measured by western blot in the cortex at 3-months of age in male and female mice. ANOVA analysis indicated that an additional copy of the Dp3Tyb region increased the abundance of **(B)** DYRK1A (F(1,49) = 16.511, p < 0.001) and **(C)** BACE2 F(1,49) = 4.444, p = 0.040). Post-hoc pair-wise comparison with Bonferroni correction for multiple comparison, demonstrated that significantly higher levels of DYRK1A were oberved in Dp3Tyb*;App^NL-F/NL-F^* compared to *App^NL-F/NL-F^* cortex (p=0.002) but that BACE2 levels did not differ between these two genotypes of mice (p= 0.359). There was no effect of an extra copy of the Dp3Tyb region on the abundance of **(E)** FL-APP level (F(1,49) = 2.183, p = 0.126), **(F)** CTF-α (F(1,40) = 0.040, p = 0.843), or **(G)** CTF-β (F(1,40) = 0.008, p = 0.929). As previously reported [15], mice homozygous for the *App^NL-F^* allele had lower abundance of **(E)** FL-APP (F(1,49) = 16.790, p < 0.001), **(F)** CTF-α (F(1,40) = 15.739, p < 0.001) and a higher abundance of **(G)** CTF-β (F(1,40) = 147.440, p < 0.001). Wild-type (WT) (female n = 6, male n = 11), Dp3Tyb (female n = 5, male n = 7), *App^NL-F^* (female n = 8, male n = 7) and Dp3Tyb*;App^NL-F/NL-F^* (female n = 5, male n = 8). CTF were below the limit of detection in Wild-type (n = 3) Dp3Tyb (n = 1), *App^NL-F^* (n = 2) and Dp3Tyb*;App^NL-F/NL-F^* (n=3) samples. Error bars show SEM, data points are independent mice. Dp3Tyb abbreviated to Dp3 for clarity.

However, we found no evidence of changes to the abundance of humanised-APP or mouse-APP in Dp3Tyb;*App^NL-F/NL-F^* or Dp3Tyb models compared to *App^NL-F/NL-F^* controls in the cortex at 3-months of age (Fig. 6D-E). Similarly no change in human-CTF-β or mouse-CTFβ was observed in the Dp3Tyb model (Fig. 6F, G). We also found no change in human/mouse APP or CTF-β abundance in the Dp(10)2Yey;*App^NL-F/NL-F^* mouse (Supplementary Fig. 5). We found no evidence of changes to total mouse amyloid-β40 or amyloid-β42 in young Dp3Tyb compared with wild-type controls (3-months of age) or insoluble human amyloid-β40 or amyloid-β42 in young (3-months of age) Dp3Tyb;*App^NL-F/NL-F^* compared with *App^NL-F/NL-F^* controls (Fig. 7, Supplementary Fig 6). These data suggest that the decreased accumulation of human amyloid-β in the Dp3Tyb model is not likely to be the result of changed abundance of APP, CTF-β or amyloid-β in the young brain and a mechanism that decreases aggregation or enhanced clearance of amyloid-β may be causal, consistent with the data from the Tc1;*App^NL-^ F/NL-F* model.

**Figure 7:**
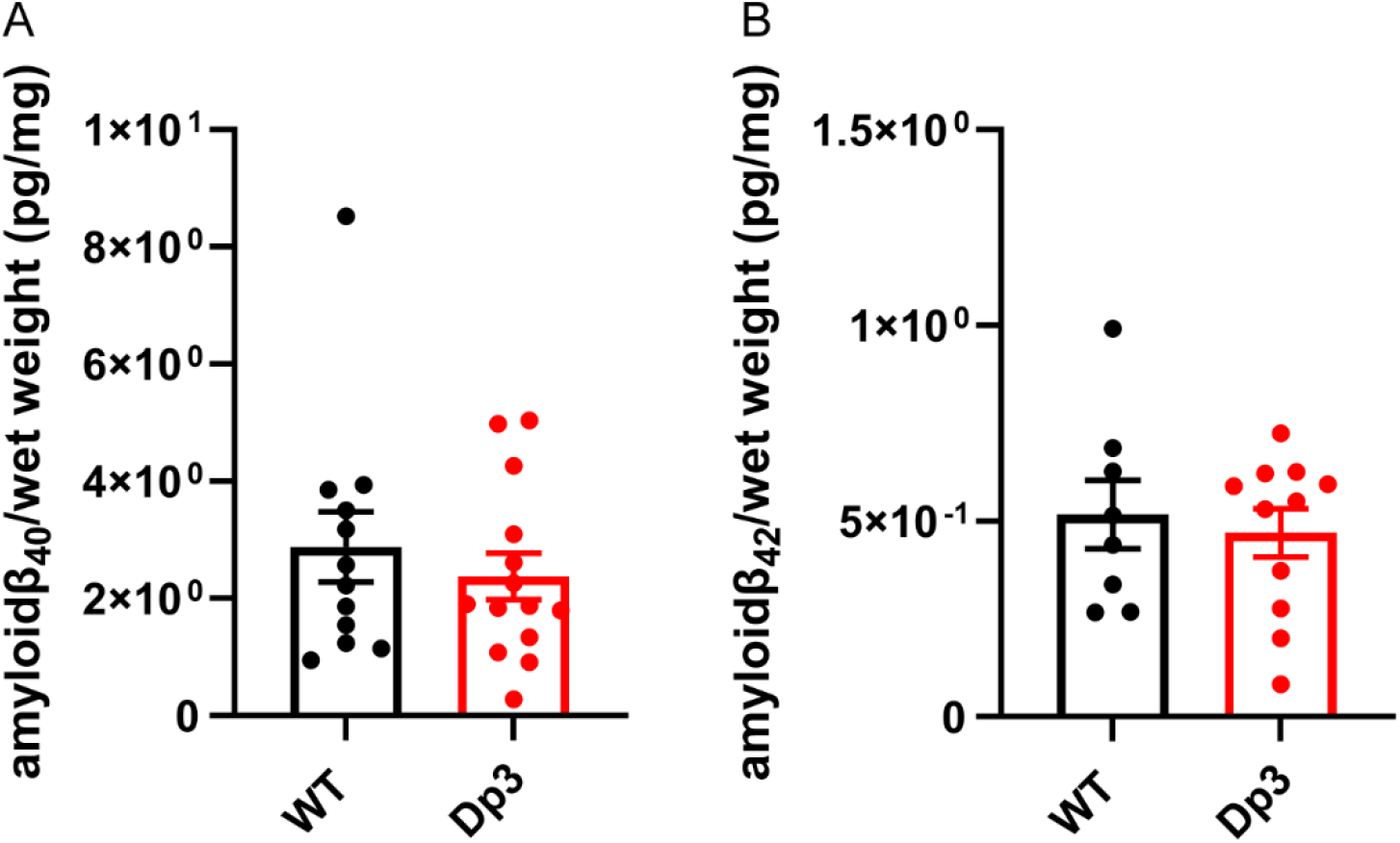
An additional copy of the Dp3Tyb region does not alter the abundance of mouse amyloid-β in the young cortex. Total cortical proteins were homogenised in TBS buffer, from mice of 3-months of age, and amyloid-β abundance was analysed by MSD assay (4G8). An additional copy of the Dp3Tyb region did not alter the abundance of **(A)** amyloid-β_40_ (F1,14) = 0.693, p = 0.419) or **(B)** amyloid-β_42_ (F1,7) = 0.410, p = 0.718). Wild-type (WT) (female n = 5, male n = 8), Dp3Tyb (female n = 7, male n = 12). Error bars show SEM, data points are independent mice. Dp3Tyb abbreviated to Dp3 for clarity.

### In Dp3Tyb mice three-copies of *Bace2* do not raise BACE2 abundance or decrease amyloid-β abundance in the young adult brain

The Dp3Tyb model carries an extra copy of *Bace2*, which encodes a secretase that has been previously reported to cleave APP at the θ site, resulting in the production of amyloid-β_1-19_ [10, 29]. BACE2 has also been suggested to clear amyloid-β, leading to reduced accumulation and production of amyloid-β_1-20_ and amyloid-β_1-34_ in an organoid model of DS [10, 29]. Thus, we wanted to investigate BACE2 in our AD-DS model. Using western blotting we found that an extra copy of the Dp3Tyb region did not cause BACE2 abundance to be significantly higher in the Dp3Tyb;*App^NL-F/NL-F^* compared with *App^NL-F/NL-F^* cortex (Fig 6A, C), likely because of the underlying high variability in the abundance of this protein in the cortex. We went on to determine if the putative BACE2 amyloid-β degradation products, human-amyloid-β_1-20_ and human-amyloid-β_1-34_ or the APP-θ cleavage product human-amyloid-β_1-19_ were altered by the Dp3Tyb region. No difference in these analytes and human-amyloid-β_1-14_, human-amyloid-β_1-15_, human-amyloid-β_1-16_ or human-amyloid-β_1-17_ was observed between Dp3Tyb;*App^NL-F/NL-F^* and *App^NL-F/NL-F^* cortex at 3-months of age (Fig. 8, Supplementary Fig. 3).

**Figure 8:**
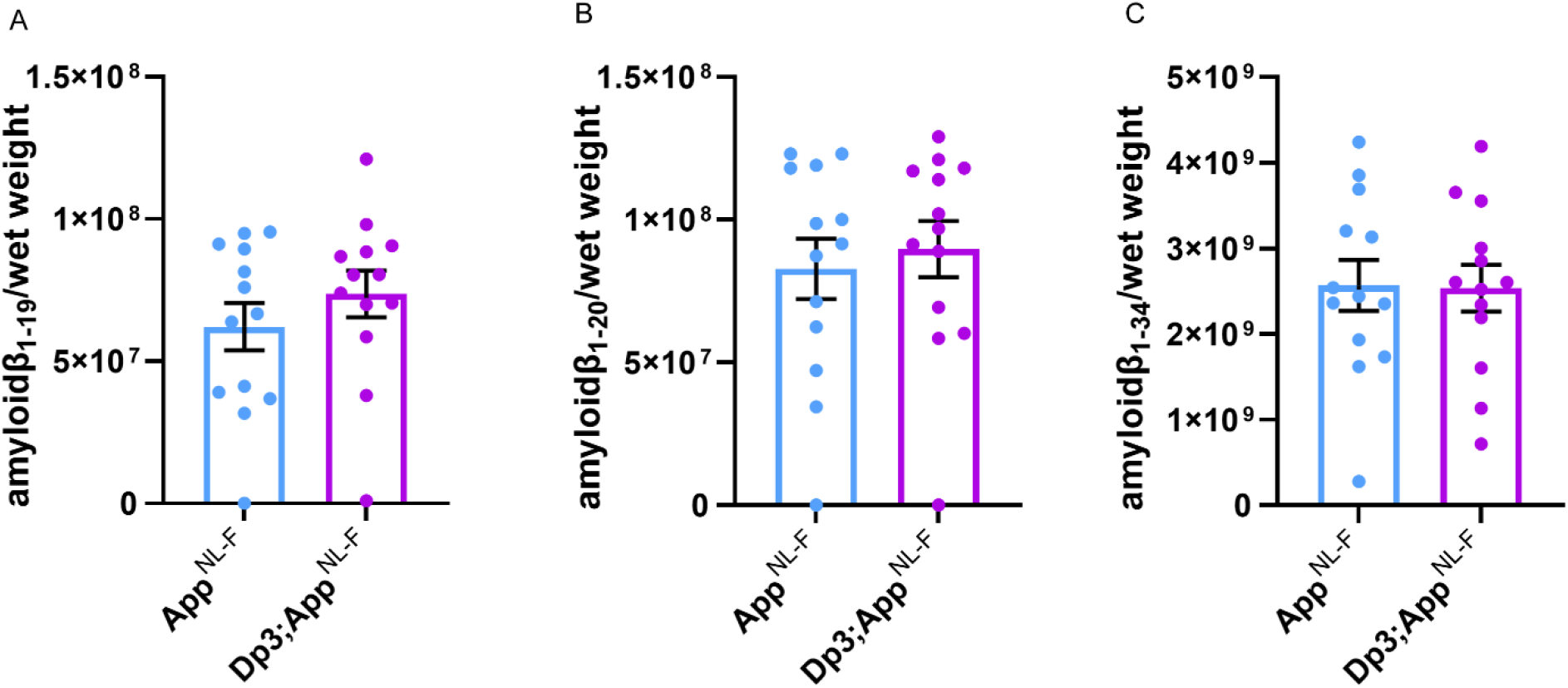
BACE2 amyloid-β cleaved fragments do not have increased abundance in the young cortex of the Dp3Tyb; *App^NL-F/NL-F^* model. LC-MS analysis of immunoprecipated cortical amyloid-β from FA fraction normalised to weight of cortical tissue, was used to determine if the Dp3Tyb region was sufficent to alter the abundance of putative BACE2 cleavage products at 3-months of age. No significant increase in the abundance of **(A)** amyloid-β1-19, (F(1,20) = 0.166 p = 0.688), **(B)** amyloid-β1-20 (F(1,20) = 0.274 p = 0.607) or **(C)** amyloid-β1-34 (F(1,20) = 0.005 p = 0.942) was observed. *App^NL-F/NL-F^* (female n = 8, male = 5), Dp3Tyb;*App^NL-F/NL-F^* (female n = 4, male = 9). Dp3Tyb abbreviated to Dp3 for clarity. Error bars show SEM, data points are independent mice. Amyloid-β1-19 and Amyloid-β1-20 n = 1 sample were below the limit of detection per genotype (these sample are shown on the graphs but were excluded from ANOVA).

As we had observed significant variability in the abundance of BACE2 in the cortex, we investigated whether BACE2 protein levels within individual mice predicted the abundance of amyloid-β_1-19_, amyloid-β_1-20_ and amyloid-β_1-34_ in the same animals. No relationships between the level of BACE2 protein and between the analytes were observed (amyloid-β_1-19_ R^2^ = 0.0001, amyloid-β_1-20_ R^2^ = 0.0079, amyloid-β1-34 R^2^ = 0.0004). These data suggest that differences in BACE2 protein abundance in the young adult mouse brain are not sufficient to cause a detectable alteration in the clearance of amyloid-β or enhanced θ-cleavage. One caveat is that at 3-months of age we cannot yet detect an alteration of the aggregation of human amyloid-β in the Dp3Tyb;*App^NL-F/NL-F^*, as detected by MSD assay after biochemical isolation in formic acid (Supplementary Fig. 6). Thus, our data could be consistent with a BACE2 aging-dependent mechanism leading to the observed decrease in amyloid-β in the Dp3Tyb;*App^NL-F/NL-F^* model at 8-months of age but further analysis is required to support this.

## Discussion

Here we use a series of mouse models of DS to identify which combination of Hsa21 genes (other than *APP*) are sufficient to modulate APP/amyloid-β. Our systematic approach identified a region of Hsa21 that contains 38 genes, which is sufficient to decrease the deposition of amyloid-β *in vivo*. This region contains two lead candidate genes *Bace2* and *Dyrk1A.*

DYRK1A is widely expressed in both the developing and adult mouse brain [30, 31] and is found in all major cell types in both the human and mouse brain [32, 33]. It is a primer kinase that can phosphorylate a large number of proteins, including APP, which has been suggested to increase the protein’s abundance [25, 34]. Inhibition of the kinase in a transgenic mouse model of AD decreased amyloid-β accumulation [34, 35]. Consistent with this, in a mouse model of DS normalisation of *Dyrk1a* gene dose leads to a decrease in APP and amyloid-β abundance [7]. In the Dp3Tyb model, which contains an additional copy of *Dyrk1a*, we observe no increase in APP abundance (endogenous mouse APP and partially humanised APP_SW_) or evidence of enhanced amyloid-β accumulation. This suggests that in our knock-in model system, 3 copies of *Dyrk1a* are not sufficient to modulate APP protein abundance or promote amyloid-β accumulation.

*BACE2* has been reported to be expressed in a subset of astrocytes and neurons in the human brain [36]. In the mouse the gene is expressed in a subset of neurons with the highest level in the CA3 and subiculum and some expression is also reported in oligodendrocytes and astrocytes lining the lateral ventricles [37]. The protein may function as a θ-secretase (decreasing CTF-β), a β-secretase (increasing CTF-β), and as amyloid-β degrading protease (decreasing amyloid-β) [29, 38]; which mechanism predominates in the human brain is not well understood. A previous study in which BACE2 was overexpressed in a wild-type APP over-expressing mouse model reported no evidence of altered amyloid-β40 or amyloid-β42 abundance in the brain [39]. Conversely, knocking-down *Bace2* in the *App^NL-G-F^* mouse model of amyloid-β accumulation has been reported to decrease CTF-β and soluble amyloid-β [40]. In contrast, reducing *BACE2* copy number in human organoids produced from individuals with DS or *APP*-duplication increased amyloid-β and triggered the formation of deposits within the model system. The authors propose this occurs because of the amyloid clearance function of BACE2, as evidenced by the raised levels of amyloid-β degradation products in trisomic compared to isogenic disomic control [10].

In the Dp3Tyb model that contains an additional copy of *Bace2*, we observe no significant change in CTF-β (endogenous mouse APP and partially humanised APP_SW_), soluble amyloid-β (endogenous mouse and humanised APP_SW_) or amyloid-β degradation products in the young adult brain. However, we observe a significant decrease in amyloid-β deposition in older mice (9-months of age) consistent with BACE2’s role as an amyloid-β degrading protease [10]. Further studies are required in aged mice to determine if enhanced amyloid-clearance can be detected or if BACE2 gene-dose correction is sufficient to reverse the reduction in amyloid accumulation caused by the Dp3Tyb model.

DS is caused by the increase in the abundance of Hsa21 gene products; however, not all genes on the chromosome are dosage-sensitive in all contexts. For example, *APP* dosage sensitivity has been reported to vary over life-span [41] and BACE2 has been reported to be both dosage-sensitive [42] and insensitive in the human brain [4, 36]. In contrast, the abundance of DYRK1A is highly sensitive to gene dose in the vast majority of reported studies [43]. Consistent with this, here we report a significant increase in DYRK1A and but could not detect a significant increase in BACE2 caused by the higher intra-animal variation in abundance of this protein in the mouse brain. Simliar overlapping levels of gene expression of *Bace2* have been previously reported at RNA level in an alternative mouse model of DS compared with euploid controls in the brain [44].

The relative abundance of proteins in AD-DS model systems is particularly important, as Down syndrome is caused by an inbalance in the relative abundance of gene products. Previous research using *APP* transgenic models, which over-express APP protein, may mask the subtle but physiologically relevant effects of the 50% increase in the abundance of other Hsa21 gene products. This study addresses this limitation, here we systematically investigate the effect of additional copies of Hsa21 gene orthologues other than *App,* on APP biology in a knock-in mouse model system. However, because of technical limitations we were not able to study the effect of an extra copy of genes located close to *App*, including the role of key genes such as *Adamts1* [45] and *Usp25* [46]. We were also unable to determine if genes located far apart on the chromosome act synergistically to cause a modulation of APP biology. Thus our data does not preclude a multigenic role for the genes in the Dp(10)2Yey or Dp(17)3Yey regions when combined with other Hsa21 genes. A further limitation is the use of AD-associated mutations in APP to drive pathology as these mutations alter the subcellular localisation and processing of APP [13] which may modulate the effect of trisomy of Hsa21 on these processes. We have partially addressed this issue by undertaking a side-by-side comparison of mouse APP/amyloid-β and partially humanised APP/amyloid-β.

## Conclusion

DS is a complex condition that alters multiple aspects of neurobiology and physiology. Here we demonstrate using physiological mouse models that an additional copy of Hsa21 genes reduces the accumulation of amyloid-β within the brain; one of the earliest steps in the AD pathogenic process. Thus trisomy of Hsa21 may partially protect individuals who have DS from the accumulation of amyloid-β, resultant from their extra copy of *APP*. Moreover we predict that individuals who have DS will accumulate amyloid-β more slowly than other groups who develop other genetic forms of early-onset AD, including that caused by duplication of *APP*. Treatments for AD-DS and other medical conditions associated with DS must take into account this fundamentally different biology; to ensure that interventions do not reverse the beneficial effect of the additional genes.

## Abbreviations

(AD): Alzheimer’s disease
(AD-DS): Alzheimer’s disease in Down syndrome
(APP): Amyloid precursor protein
(BSA): Bovine serum albumin
(DS): Down syndrome
(LC-MS): Liquid chromatography-mass spectrometry
(MSD): Meso Scale Discovery
(PBS): Phosphate buffered saline
(SEM): Standard error of the mean

## Author Contributions

P.M. undertook biochemical and histological experiments, data analysis and edited the manuscript. J.T and S.A undertook biochemical and histological experiments. G.L undertook histological analysis. S.N undertook histological experiments and K.C. undertook genotyping. E.G.W. and G.B. performed the LC-MS analysis. E.Y., T. S. and T.C.S. contributed essential research resources, E.M.C.F. and V.L.J.T, contributed essential research resources, designed and supervised the study and edited the manuscript. F.K.W. designed and supervised the study, undertook data analysis and wrote the manuscript. All authors revised the manuscript.

## Acknowledgements

For the purpose of Open Access, the author has applied a CC-BY public copyright licence to any Author Accepted Manuscript version arising from this submission. All authors read and approved the final manuscript. We thank Dr. Amanda Heslegrave (UCL) for assistance with this project.

