## Supplementary Information for "Genetic mapping of APP and amyloid-β biology modulation by trisomy 21"

**Supplementary Data**

Paige Mumford[1,*], Justin Tosh[2,*], Silvia Anderle[1], Eleni Gkanatsiou Wikberg[3], Gloria Lau[1], Sue Noy[2], Karen Cleverley[2], Takashi Saito[4], Takaomi C Saido[4], Eugene Y. Yu[5], Gunnar Brinkmalm[3], Erik Portelius[3], Kaj Blennow[3,6], Henrik Zetterberg[1,3,6,7,8], Victor Tybulewicz[9,10, 11], Elizabeth M.C. Fisher¶[2,11] and Frances K. Wiseman¶[1,11]

*These authors contributed equally

¶Corresponding authors.

 (E.M.C.F.) or (F.K.W.)

**Supplementary Figure 1**

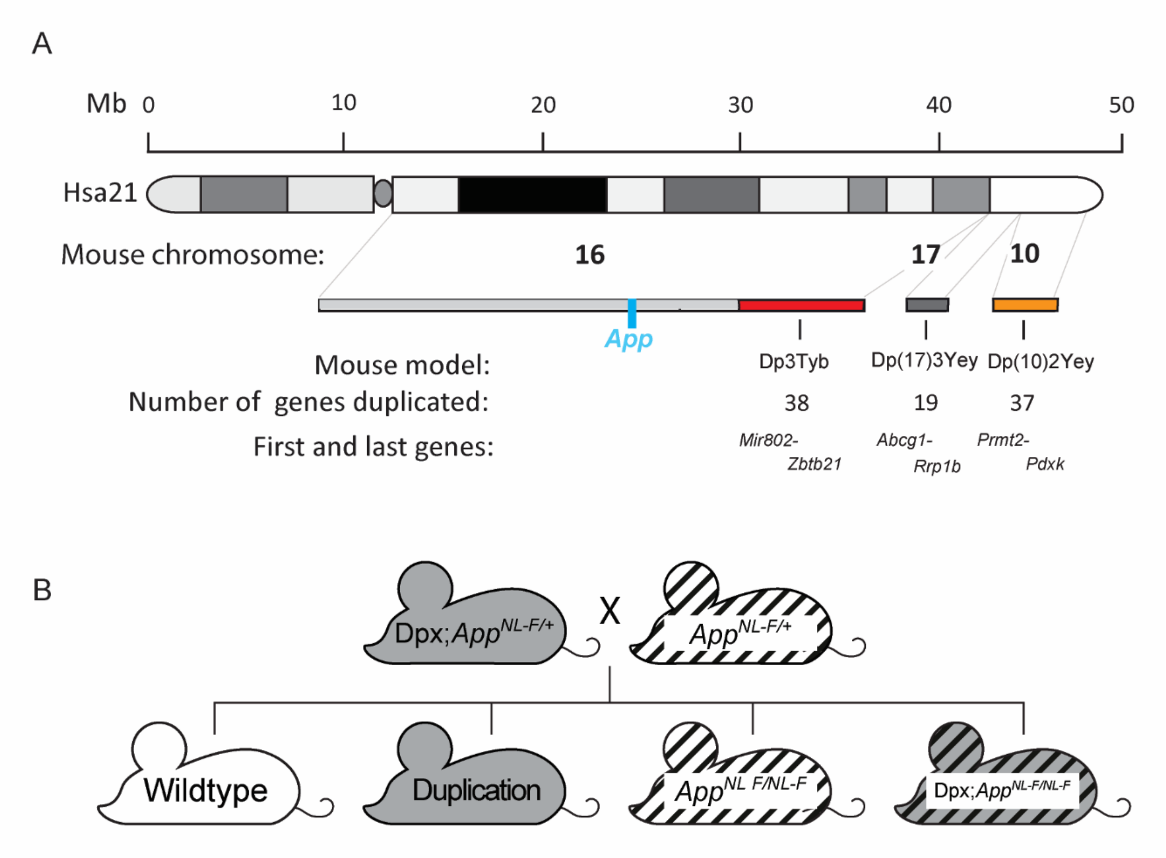


**(A)** Schematic of the genetic content of the segmental duplication mouse models (Dp3Tyb, Dp(10)2Yey and Dp(17)3Yey). (**B)** Schematic of the generation of experimental cohorts for the duplication models (Dp(10)2Yey, Dp(17)3Yey and Dp3Tyb) with the studies were produced by crossing mice carrying the duplications Dpx;*App*$^{NL\text{-}F/+}$ with *App*$^{NL\text{-}F/+}$ mice**.**

**Supplementary Figure 2**

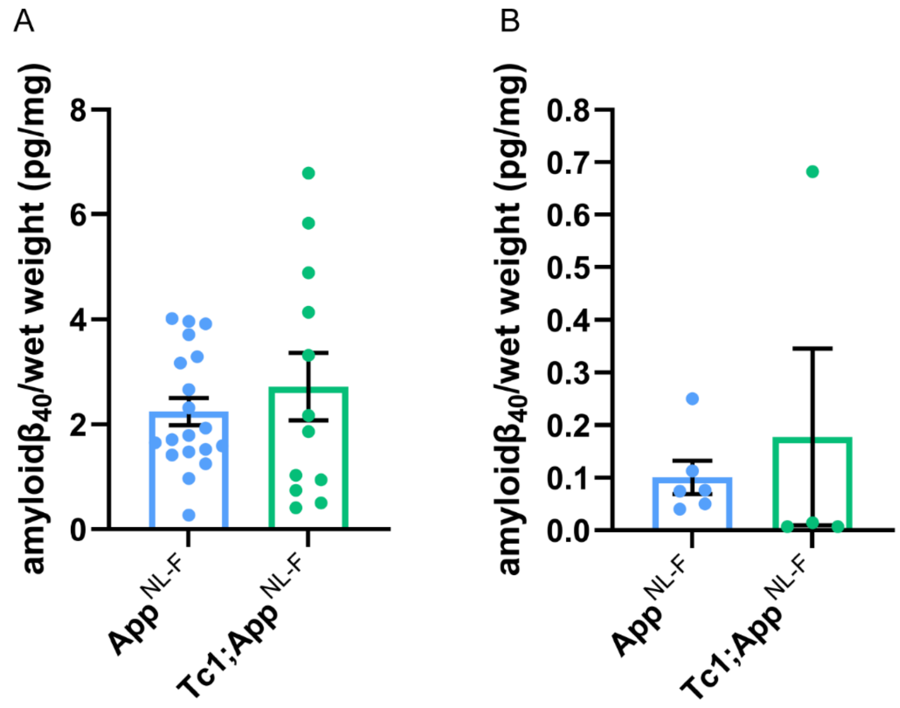


Total cortical proteins were biochemically fractionated from mice of 8-months of age and amyloid abundance analysed by MSD assay. Error bars show SEM, data points are independent mice. **(A)** No difference in the abundance of 5 M guanidine hydrochloride soluble amyloid-$\beta_{40}$ was observed between Tc1;*App*$^{NL-F/NL-F}$ compared with *App*$^{NL-F/NL-F}$ controls ($F(1,13) = 0.005$, $p = 0.946$). *App*$^{NL-F/NL-F}$ female n = 11, male n = 8, n = 4 below limit of detection; Tc1;*App*$^{NL-F/NL-F}$ female n = 7, male n = 5, n = 6 below limit of detection. **(B)** No difference in the abundance of 1% Triton X-100 soluble amyloid-$\beta_{40}$ was observed between Tc1;*App*$^{NL-F/NL-F}$ compared with *App*$^{NL-F/NL-F}$ controld ($F(1,3) = 0.000$, $p = 0.991$). *App*$^{NL-F/NL-F}$ female n = 4, male n = 0 (n = 17 below limit of detection); Tc1;*App*$^{NL-F/NL-F}$ female n = 4, male n = 2 (n = 14 below limit of detection). Tris soluble amyloid-$\beta_{40}$ was not detected in these samples.

**Supplementary Figure 3**

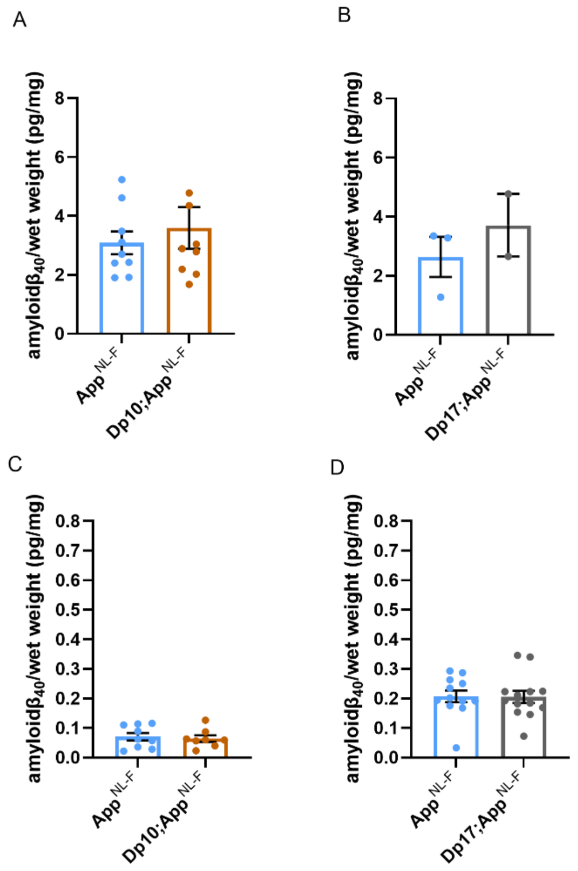


Total cortical proteins were biochemically fractionated from mice of 8-months of age and amyloid abundance analysed by MSD assay. **(A)** No difference in the abundance of 5 M guanidine hydrochloride soluble amyloid-$\beta_{40}$ was observed between Dp(10)2Yey;*App*$^{NL-F/NL-F}$ compared with *App*$^{NL-F/NL-F}$ controls ($F(1,12) = 2.137$ , $p = 0.169$). *App*$^{NL-F/NL-F}$ female n = 6, male n = 3; Dp(10)2Yey*;App*$^{NL-F/NL-F}$ female n = 4, male n = 5. **(B)** No difference in the abundance of 5 M guanidine hydrochloride soluble amyloid-$\beta_{40}$ was observed between Dp(17)3Yey;*App*$^{NL-F/NL-F}$ compared with *App*$^{NL-F/NL-F}$ controls ($F(1,1) = 0.782$, $p = 0.539$). *App*$^{NL-F/NL-F}$ female n = 1, male n = 2, n = 9 below limit of detection; Dp(17)3Yey*;App*$^{NL-F/NL-F}$ female n = 2, male n = 0, n = 12 below limit of detection. **(C)** No difference in the abundance of 1% Triton X-100 soluble amyloid-$\beta_{40}$ was observed between Dp(10)2Yey;*App*$^{NL-F/NL-F}$ compared with *App*$^{NL-F/NL-F}$ controls ($F(1,11) = 0.540$, $p = 0.478$). *App*$^{NL-F/NL-F}$ female n = 6, male n = 3; Dp(10)2Yey*;App*$^{NL-F/NL-F}$ female n = 4, male n = 4. **(D)** No difference in the abundance of 1% Triton X-100 soluble amyloid-$\beta_{40}$ was observed between Dp(17)3Yey;*App*$^{NL-F/NL-F}$ compared with *App*$^{NL-F/NL-F}$ controls ($F(1,19) = 0.007$, $p = 0.982$). *App*$^{NL-F/NL-F}$ female n = 7, male n = 5; Dp(10)2Yey*;App*$^{NL-F/NL-F}$ female n = 7, male n = 6, n = 1 below limit of detection. Error bars show SEM, data points are independent mice. Dp(10)2Yey abbreviated to Dp10 and Dp(17)3Yey abbreviated to Dp17 for clarity.

**Supplementary Figure 4**

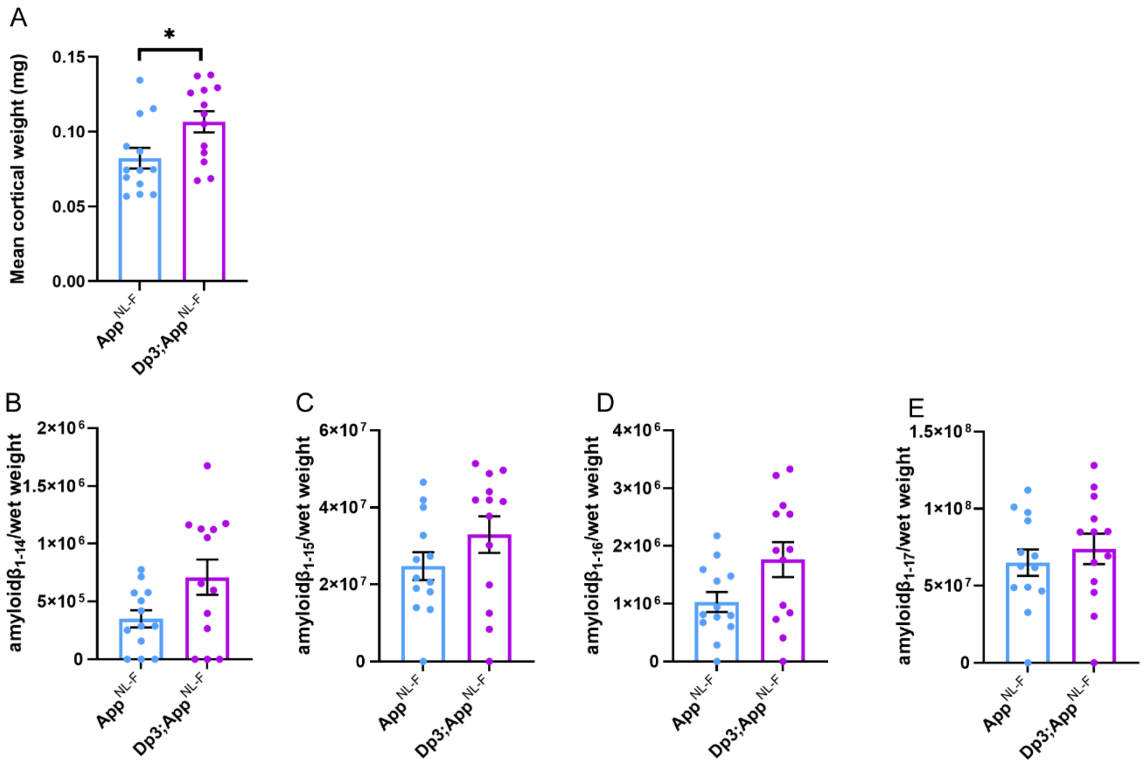


**(A)** Dp3Tyb;*App*$^{NL\text{-}F/NL\text{-}F}$ cortex weighs more than *App*$^{NL\text{-}F/NL\text{-}F}$ cortex at 3-months of age ($F(1,22)= 7.772$, $p=0.011$). *App*$^{NL\text{-}F/NL\text{-}F}$ (female n = 8, male = 5), Dp3Tyb;*App*$^{NL\text{-}F/NL\text{-}F}$ (female n = 4, male = 9). **(B)-(E)** LC-MS analysis of immunoprecipitated cortical amyloid-β from FA fraction normalised to weight of cortical tissue at 3-months of age. No difference in the abundance of **(B)** amyloid-$\beta_{1\text{-}14,}$ ($F(1,15) = 1.622$, $p = 0.222$), **(C)** amyloid-$\beta_{1\text{-}15,}$ ($F(1,19) = 0.496$, $p = 0.490$), **(D)** amyloid-$\beta_{1\text{-}16,}$ ($F(1,19) = 2.274$, $p = 0.148$) or **(E)** amyloid-$\beta_{1\text{-}17,}$ ($F(1,19) = 0.079$, $p = 0.781$) was detected in the cortex of *App*$^{NL\text{-}F/NL\text{-}F}$ compared with *App*$^{NL\text{-}F/NL\text{-}F}$ mice. Amyloid-$\beta_{1\text{-}14}$ n = 3 samples were below the limit of detection per genotype, amyloid-$\beta_{1\text{-}15,\ 1\text{-}16,\ 1\text{-}17}$ n = 1 samples were below the limit of detection per genotype (these sample are shown on the graphs but were excluded from ANOVA). Error bars show SEM, data points are independent mice. Dp3Tyb abbreviated to Dp3 for clarity.

**Supplementary Figure 5**

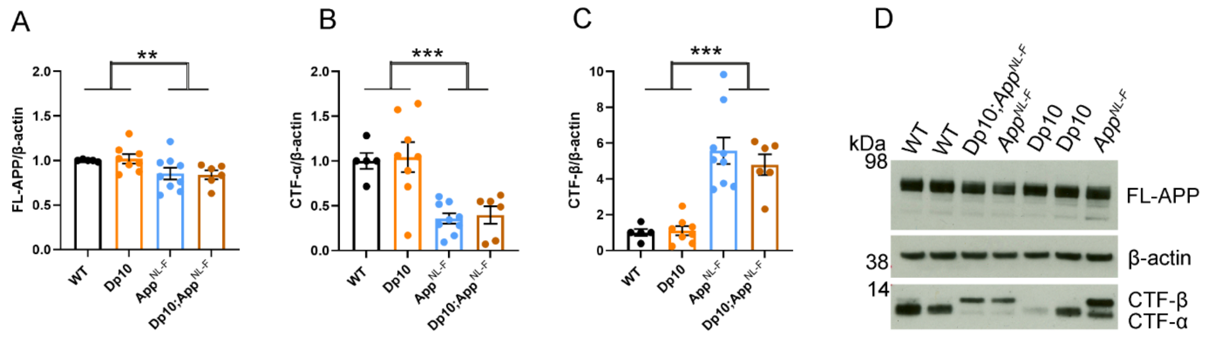


The abundance of **(A, D)** full-length, APP (FL-APP), **(B, D)** C-terminal fragment-α (CTF-α) and **(C, D)** C-terminal fragment-β (CTF-β) relative to β-actin loading control was measured by western blot in the cortex at 3-months of age in male and female mice. There was no effect of an extra copy of the Dp(10)2Yey region on the abundance of **(A)** FL-APP level ($F(1,22) = 0.828$, $p = 0.372$), **(B)** CTF-α ($F(1,22) = 0.054$, $p = 0.819$) or **(C)** CTF-β abundance ($F(1,22) = 0.829$, $p = 0.372$). As previously observed, mice homozygous for the $App^{NL\text{-}F}$ allele had lower abundance of **(A)** FL-APP ($F(1,22) = 8.168$, $p = 0.009$), **(B)** CTF-α ($F(1,22) = 72.150$, $p < 0.001$) and a higher abundance of **(C)** CTF-β ($F(1,22) = 15.300$, $p < 0.001$). Wild-type (WT) (female n = 1, male n = 4), Dp3Tyb (male n = 8), $App^{NL\text{-}F}$ (male n = 9) and Dp3Tyb;$App^{NL\text{-}F/NL\text{-}F}$ (female n = 1, male n = 4). Error bars show SEM, data points are independent mice. Dp(10)2Yey abbreviated to Dp10 for clarity.

**Supplementary Figure 6**

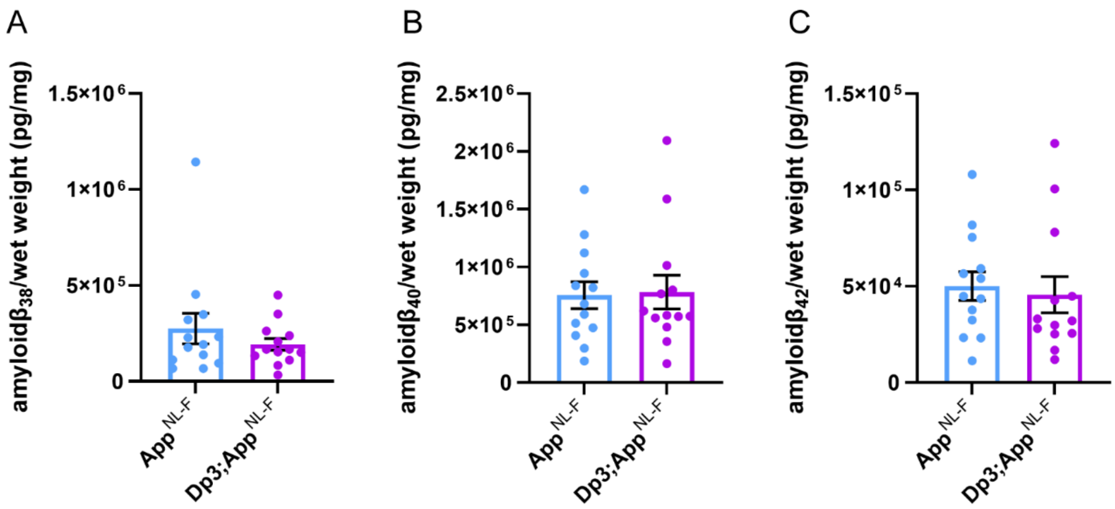


Total cortical proteins of 3-month old mice were prepared for mass-spectrometry analysis and amyloid-β abundance analysed by MSD assay (6E10). No difference in the abundance of formic acid soluble **(A)** amyloid-$\beta_{38}$ (F(1,21) = 1.001, p = 0.328), **(B)** amyloid-$\beta_{40}$ (F(1,21) = 0.032 p = 0.860) or **(C)** amyloid-$\beta_{42}$ (F(1,21) = 0.306, p = 0.443) was detected. *App*$^{NL\text{-}F/NL\text{-}F}$ (female n = 8, male = 5), Dp3Tyb;*App*$^{NL\text{-}F/NL\text{-}F}$ (female n = 4, male = 9). Dp3Tyb abbreviated to Dp3 for clarity. Error bars show SEM, data points are independent mice.
